# Signaling through FcγRIIA and the C5a-C5aR pathway mediates platelet hyperactivation in COVID-19

**DOI:** 10.1101/2021.05.01.442279

**Authors:** Sokratis A. Apostolidis, Amrita Sarkar, Heather M. Giannini, Rishi R. Goel, Divij Mathew, Aae Suzuki, Amy E. Baxter, Allison R. Greenplate, Cécile Alanio, Mohamed Abdel-Hakeem, Derek A. Oldridge, Josephine Giles, Jennifer E. Wu, Zeyu Chen, Yinghui Jane Huang, Ajinkya Pattekar, Sasikanth Manne, Oliva Kuthuru, Jeanette Dougherty, Brittany Weiderhold, Ariel R. Weisman, Caroline A. G. Ittner, Sigrid Gouma, Debora Dunbar, Ian Frank, Alexander C. Huang, Laura A. Vella, The UPenn COVID Processing Unit, John P. Reilly, Scott E. Hensley, Lubica Rauova, Liang Zhao, Nuala J. Meyer, Mortimer Poncz, Charles S. Abrams, E. John Wherry

## Abstract

Patients with COVID-19 present with a wide variety of clinical manifestations. Thromboembolic events constitute a significant cause of morbidity and mortality in patients infected with SARS-CoV-2. Severe COVID-19 has been associated with hyperinflammation and pre-existing cardiovascular disease. Platelets are important mediators and sensors of inflammation and are directly affected by cardiovascular stressors. In this report, we found that platelets from severely ill, hospitalized COVID-19 patients exhibit higher basal levels of activation measured by P-selectin surface expression, and have a poor functional reserve upon *in vitro* stimulation. Correlating clinical features to the ability of plasma from COVID-19 patients to stimulate control platelets identified ferritin as a pivotal clinical marker associated with platelet hyperactivation. The COVID-19 plasma-mediated effect on control platelets was highest for patients that subsequently developed inpatient thrombotic events. Proteomic analysis of plasma from COVID-19 patients identified key mediators of inflammation and cardiovascular disease that positively correlated with *in vitro* platelet activation. Mechanistically, blocking the signaling of the FcγRIIa-Syk and C5a-C5aR pathways on platelets, using antibody-mediated neutralization, IgG depletion or the Syk inhibitor fostamatinib, reversed this hyperactivity driven by COVID-19 plasma and prevented platelet aggregation in endothelial microfluidic chamber conditions, thus identifying these potentially actionable pathways as central for platelet activation and/or vascular complications in COVID-19 patients. In conclusion, we reveal a key role of platelet-mediated immunothrombosis in COVID-19 and identify distinct, clinically relevant, targetable signaling pathways that mediate this effect. These studies have implications for the role of platelet hyperactivation in complications associated with SARS-CoV-2 infection.

**Cover illustration:** 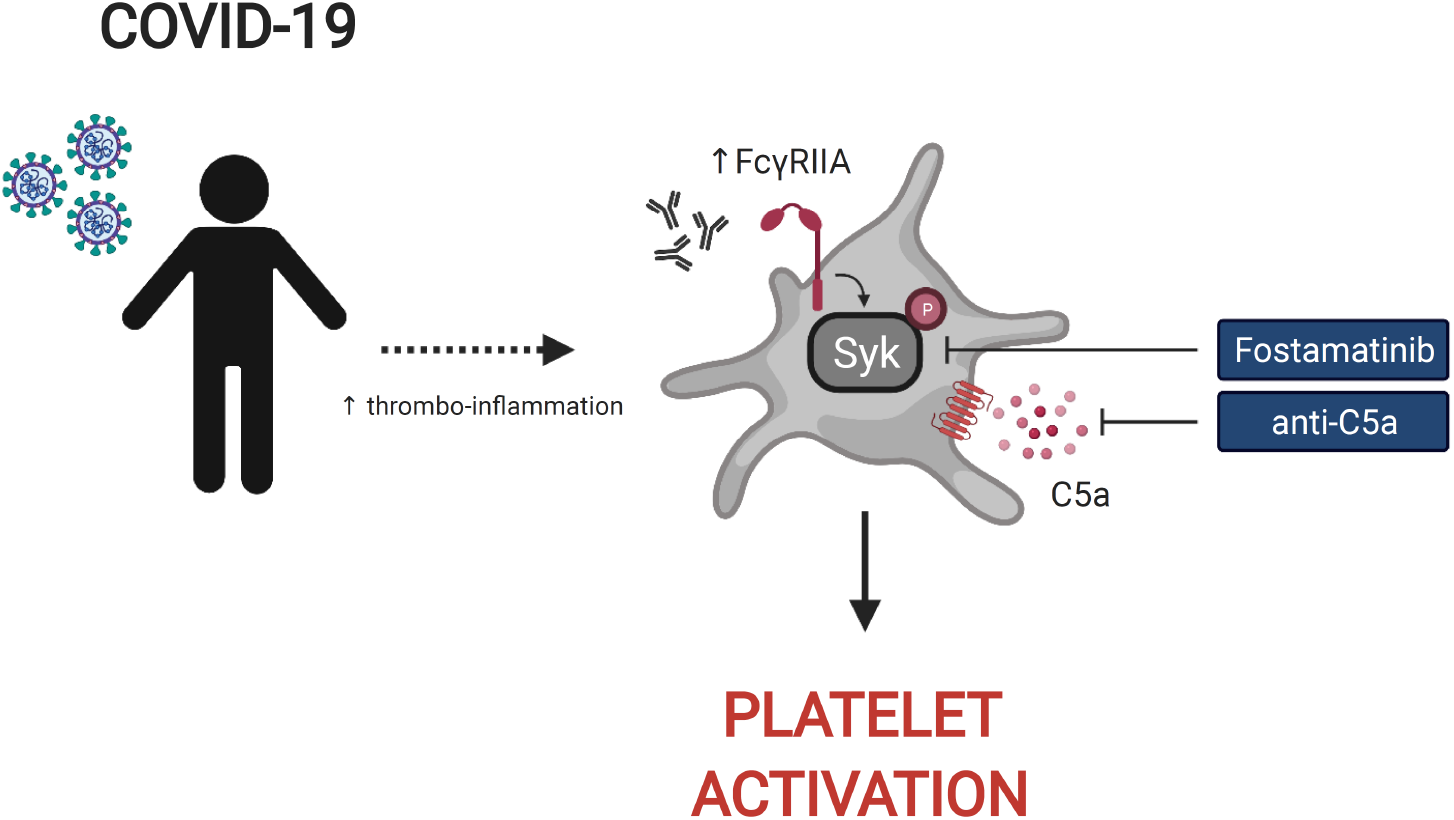

**One-sentence summary:** The FcγRIIA and C5a-C5aR pathways mediate platelet hyperactivation in COVID-19

## Introduction

Coronavirus disease 2019 (COVID-19) has led to a global-scale pandemic creating an unprecedented burden on human health and public health processes (*1*). SARS-CoV-2 infected patients represent a wide spectrum of clinical presentations, ranging from asymptomatic infections to prolonged ICU stays accompanied by significant morbidity and mortality (*2-4*). Although Acute Respiratory Distress Syndrome (ARDS) represents the hallmark of COVID-19 associated clinical manifestations, thrombotic events are enriched in patients with severe COVID-19 and have been linked to worse outcomes (*3, 5, 6*). Increased levels of d-dimer and platelet dysfunction are frequently observed in COVID-19 patients (*7–11*), indicating a loss of homeostasis in platelet function, vascular integrity and the coagulation cascade.

Platelets are anucleated megakaryocyte-derived blood cells that play a prominent role in hemostasis and thrombus formation (*12*). Beyond hemostasis, platelets represent cellular mediators of inflammation and interact with the immune system in multiple ways, including priming of other immune cells and integrating extrinsic immunological stimuli (*13–15*). Platelets express a variety of TLR receptors, express HLA class II for antigen presentation and respond to complement activation (*13, 16, 17*). In COVID-19, patients with severe disease often exhibit increased platelet activation and formation of platelet-monocyte aggregates facilitated by tissue factor expression on monocytes (*18*). RNA sequencing of platelets in COVID-19 patients has revealed an altered transcriptional profile with enrichment in the pathways of antigen presentation, protein ubiquitination and mitochondrial dysfunction (*10*). A candidate-driven genetic association study identified putative complement and coagulation-related loci associated with severe COVID-19 (*19*). Finally, unbiased pathway-enrichment analysis of circulating proteins in COVID-19 patients underscored platelet degranulation and complement activation as the top pathways associated with disease severity (*20*). Thus, platelets have key role not only in hemostasis, but also in inflammatory processes and platelet dysregulation is central to the pathogenesis of severe COVID-19 in many patients.

Despite the importance of platelets in thrombotic events in COVID-19 patients, how heightened platelet activation is linked to clinical features of disease, and the associated underlying mechanisms remain poorly understood. These gaps in our understanding of platelet function and dysfunction during SARS-CoV-2 infection limit our ability to identify patients at risk of thromboembolic events and to treat vascular complications of COVID-19 including clots. Moreover, identifying the inflammatory effector molecules and pathways that underlie the activation and dysregulation of platelets in COVID-19 could reveal novel opportunities for therapeutic intervention. To address these questions, we examined platelets and the platelet-activating potential of plasma from severely ill, hospitalized COVID-19 patients. These studies revealed an increase in basal expression of the activation marker P-selectin on platelets from severe COVID-19 patients coupled with poor response to TRAP stimulation, indicating loss of functional reserve, compared to platelets from convalescent and healthy donors. COVID-19 patient plasma robustly activated healthy platelets from control donors and this platelet activating potential was highest prior to the precipitation of a thrombotic event. Correlation of platelet activation induced by COVID-19 plasma with clinical features collected during patient hospitalization revealed significant associations with ferritin levels. Moreover, proteomic analysis identified central mediators of inflammation and cardiovascular homeostasis correlating with a platelet hyperactivated state consistent with a role for platelets linking inflammatory events to thrombotic pathology. Finally, we identified a key role for Fc receptor and complement signaling in platelet activation in COVID-19 because blockade of signaling through the FcγRIIa-Syk and the C5a-C5aR axis using antibody blockade, depletion of immunoglobulin from COVID-19 plasma or the FDA approved drug fostamatinib blocked activation of healthy platelets by COVID-19 plasma. Thus, these studies identify a platelet hyperactive state associated with severe SARS-CoV-2 infection, define the underlying mechanisms and have direct therapeutic implications for the prevention and treatment of thrombotic complications in patients with COVID-19.

## Results

### Platelets from hospitalized COVID-19 patients exhibit increased activation at baseline and poor functional reserve

COVID-19 is associated with heightened activation of platelets both at baseline and after pharmacologic stimulation (*18, 21*). We interrogated a cohort of hospitalized COVID-19 patients, non-hospitalized COVID-19 recovered subjects and healthy control subjects recruited at the University of Pennsylvania for which we had collected peripheral blood samples and clinical annotation (**Supplementary Table 1**) (*22*). The inpatient cohort spanned a range of severity scores with moderate and severe/critical scores being the most common. Most patients were treated in a high-acuity medicine floor or ICU setting and clinically manifested pneumonia with hypoxia.

CD62P (P-selectin) is bound to the membrane of α-granules within the cytoplasm of platelets, transported to the plasma membrane rapidly after activation and can be used as a marker of platelet activation. To evaluate the activation status of platelets from COVID-19 patients in our cohort, we assayed surface expression of CD62P directly *ex vivo* or after stimulation with Thrombin Receptor Activation Peptide (TRAP) that activates platelets via the thrombin receptor (**Figure 1A**). We compared CD62P surface expression in three separate groups: hospitalized COVID-19 patients (COVID-19 inpatient), patients previously infected with SARS-CoV-2 that recovered and reached convalescence (COVID-19 convalescent) and healthy donors. The COVID-19 inpatient group had higher expression of CD62P at baseline [median geometric mean fluorescence intensity (gMFI) 56.0] compared to the COVID-19 convalescent (median gMFI 45.2) or healthy donor (median gMFI 20.2) group (**Figure 1A** and **1B**). Upon TRAP stimulation, CD62P expression increased for all three groups (**Figure 1A** and **1B**). However, the stimulation-induced degranulation and upregulation of surface CD62P was higher for the healthy donor group compared to the other two groups (**Figure 1B**). As a result, the ratio of TRAP/basal surface CD62P expression was highest for the healthy donor group (ratio 28.4), intermediate for the convalescent group (ratio 6.98) and lowest for the inpatient COVID19 group (ratio 5.77, **Figure 1B**). These results identify a heightened basal platelet activation state in hospitalized COVID-19 patients, but also demonstrate a reduced functional reserve in platelets from these patients revealed by *ex vivo* TRAP stimulation.

**Figure 1.**
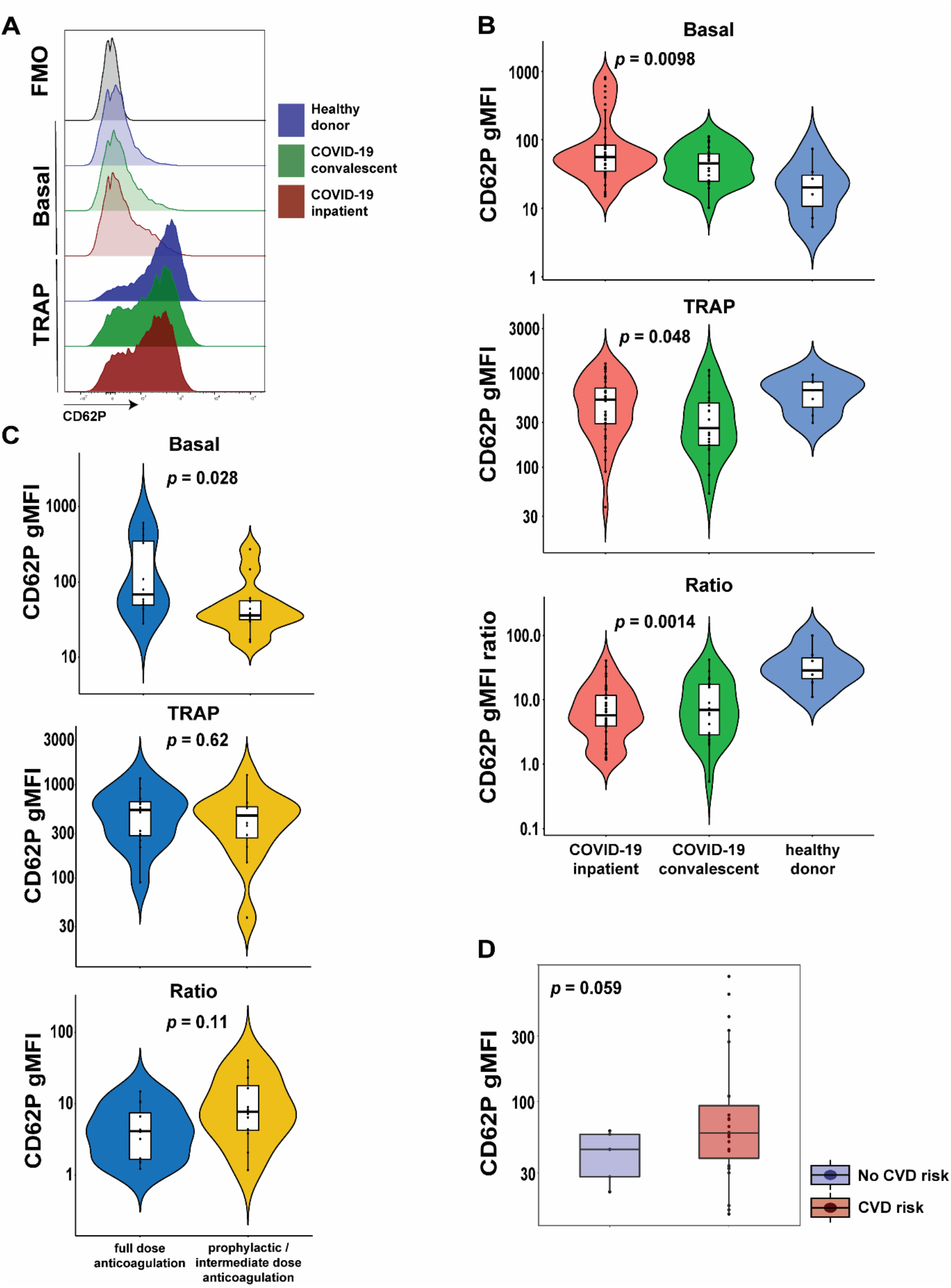
Platelets from hospitalized COVID-19 patients exhibit increased activation at baseline and diminished functional reserve. **A-B)** Representative histograms (A) and cumulative data (B) for CD62P surface expression of *ex vivo* isolated platelets assayed with or without TRAP stimulation for 20mins. The CD62P gMFI ratio of basal/TRAP-treated is also shown. The three patient cohorts shown are COVID-19 inpatient, COVID-19 convalescent and healthy donors. Kruskal-Wallis non-parametric testing was used and the *p*-values are depicted. **C)** Cumulative data for CD62P surface expression of *ex vivo* isolated platelets at baseline (basal), after TRAP activation (TRAP) and their ratio for hospitalized COVID-19 patients on full-dose or prophylactic/intermediate dose of anticoagulation. Mann-Whitney nonparametric testing was used and the *p*-values are depicted. **D)** Cumulative data for CD62P surface expression of *ex vivo* isolated platelets at baseline for hospitalized COVID-19 patients with or without cardiovascular disease risk factors. Mann-Whitney non-parametric testing was used, and the *p*-value is depicted. FMO control: Fluorescence minus one control, gMFI: geometric mean fluorescence intensity, TRAP: Thrombin Receptor Activation Peptide.

A causal link between cardiovascular disease and COVID-19 outcomes, including thrombotic complications, has been proposed previously (*23–25*) but how exactly such clinical events relate to platelet activation phenotypes is not clear. Thus, we first assessed CD62P expression, both basal and following TRAP stimulation, in hospitalized COVID-19 patients that exhibited clinically evident thrombosis. However, CD62P surface levels were similar between COVID-19 inpatients that had a clinical thrombus and those who did not (**Supplementary Figure 1**). COVID-19 hospitalized patients are placed on different protocols of anticoagulation. In our cohort, several patients were on full-dose anticoagulation and others were on standard-of-care, prophylactic dose (or intermediate according to the inpatient protocol at the time of COVID-19 hospital care) anticoagulation. This distinction likely captures both the effect of inpatient clotting incidents and pre-existing, underlying comorbidities, including pro-thrombotic states and cardiovascular etiologies, requiring anti-coagulation. We therefore used this distinction in anticoagulant treatment to evaluate the CD62P results. This analysis revealed a significant increase in basal CD62P surface levels in patients on full-dose compared to those on prophylactic or intermediate dose anticoagulation (**Figure 1C**). Both groups, however, responded after TRAP stimulation by increasing CD62P MFI and although there was a trend for lower CD62P TRAP/basal ratio in the full-dose anticoagulation group, this difference did not reach statistical significance (**Figure 1C**). To address this question from a different perspective, we compared the platelet activation indicated by CD62P expression between subjects with and without cardiovascular disease (CVD) risk factors. Indeed, COVID-19 patients with CVD risk displayed elevated basal CD62P expression on the plasma membrane of platelets compared to patients without CVD risks (**Figure 1D**). Of note, there was no association between platelet *ex vivo* surface CD62P levels and COVID-19 NIH disease severity score (Kruskal-Wallis, *p* = 0.16 for basal levels and *p* = 0.31 for TRAP/basal ratio). These data indicate that platelet activation in COVID-19 patients was higher in those patients on full-dose anticoagulation either due to an acquired in-hospital event or a pre-existing condition and was also associated with the presence of cardiovascular risk. Thus, defining the mechanisms and pathways contributing to the platelet activation may reveal insights into COVID-19 pathogenesis and thrombotic complications.

### Plasma from hospitalized COVID-19 patients with high levels of inflammatory markers causes platelet hyperactivation

COVID-19 can often manifest as a hyperinflammatory state. We, therefore, next investigated whether plasma from COVID-19 patients contained soluble mediators capable of activating platelets. To test this idea, we incubated plasma from COVID-19 patients with platelets from healthy donors and assessed platelet activation state. In addition to CD62P, we measured surface CD63 levels (also known as LAMP-3), a component of dense granules in platelet cytoplasm that translocates to the plasma membrane upon activation and subsequent degranulation. We also quantified Fcγ receptor IIa (FcγRIIa, CD32), the only Fcγ receptor on platelets, and C3aR on platelets to evaluate changes in these receptors that could underlie potential effects of immune complexes and/or anaphylatoxins on platelet activation. The control platelets were pre-gated as CD42b+ (GPIba+) single cells (for gating strategy see **Supplementary Figure 2**). Unlike our *ex vivo* analysis of platelets from COVID-19 patients, treatment of healthy platelets using plasma from COVID-19 patients did not induce higher CD62P expression than plasma from recovered or healthy donors (**Figure 2A**). In contrast, however, CD63 and CD32 were increased on control platelets following incubation with plasma from the COVID-19 inpatient group compared to platelets treated with plasma from the COVID-19 convalescent group (**Figure 2A**). C3aR levels also increased upon treatment with COVID-19 inpatient plasma when compared to the healthy donor plasma (**Figure 2A**). The reasons for the differences in CD62P between direct *ex vivo* analysis of platelets from COVID-19 patients and induction of healthy platelet activation from plasma from COVID-19 patients may reflect a different kinetics after *in vivo* versus *in vitro* activation, or non-plasma-based activation signals for CD62P. Nevertheless, these results demonstrate that plasma from COVID-19 patients preferentially activates platelets indicating that this assay can be used to further gain insights into pathways underlying platelet activation and/or dysfunction associated with COVID-19 disease.

**Figure 2.**
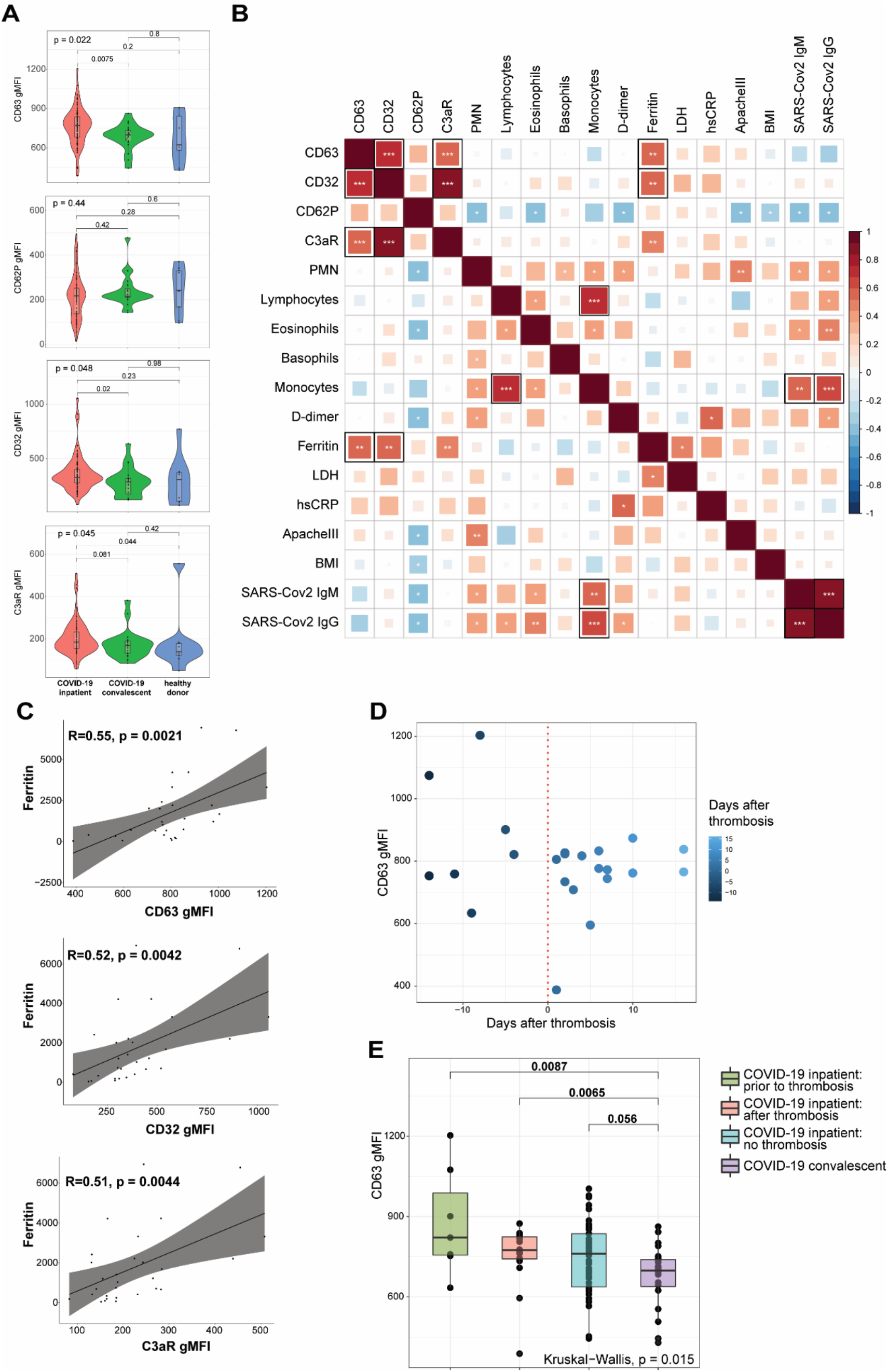
The ability of COVID19 plasma to activate platelets is increased for patients with high circulating levels of ferritin and at timepoints preceding a thrombotic event. **A)** Plasma from COVID-19 inpatient, COVID-19 convalescent and healthy donors was incubated with platelets isolated from healthy volunteers. The gMFI levels of CD63, CD62P, CD32 and C3aR are shown in these three groups. Kruskal-Wallis non-parametric testing was used to compare all three groups and pair-wise comparisons were also performed; *p*-values are depicted. **B)** Spearman correlation of the gMFI surface levels of the markers used in (A) and selected clinical parameters of the COVID-19 inpatient group. ρ correlation coefficient shown on the key (right). Asterisks *, ** and *** denote p values less than 0.05, 0.01 and 0.001 respectively. Highlighted squares denote FDR values less than 0.05. **C)** Representative scatter plots for ferritin with CD63, CD32 and C3aR. **D)** CD63 gMFI surface levels of control platelets induced by plasma derived from COVID-19 patients with thrombosis drawn at different timepoints relative to the thrombotic event. **E)** CD63 gMFI surface levels of control platelets induced by plasma derived from COVID-19 patients prior to thrombosis and after thrombosis, COVID-19 patients without thrombosis and COVID-19 patients in convalescence.

The data in **Figure 2A** implicate factors circulating in the plasma from COVID-19 patients that mediate platelet activation. However, there was considerable within-group heterogeneity. We hypothesized that this variation in platelet activation could be explained at least partially by the clinical heterogeneity documented in COVID-19 patients (*26–28*). To test this idea, we correlated surface expression of CD62P, CD63, CD32 and C3aR with clinical variables obtained during the hospitalization of COVID-19 patients (**Figure 2B**). These clinically measured features of disease included cell counts (polymorphonuclear cells, lymphocytes, eosinophils, basophils, monocytes), levels of d-dimer, ferritin, lactate dehydrogenase (LDH), high-sensitivity CRP (hs-CRP), Apache III scoring, Body-Mass index (BMI) and concentration of SARS-CoV-2 specific IgM and IgG. Analyzing these data together with platelet activation changes induced by plasma from COVID-19 patients revealed strong correlations among CD63, CD32 and C3aR, but not with CD62P (**Figure 2B**, top left 4×4 square of correlations). Examining clinical features, ferritin had the strongest positive correlation with the ability of plasma to activate platelets (**Figure 2B-C**). There was a statistically significant positive correlation between ferritin and CD63 or CD32 that also met a false discovery rate (FDR) cutoff of 0.05 (**Figure 2B**). C3aR was also positively correlated with ferritin (*p* <0.01) but this relationship did not achieve FDR <0.05. Other correlations between clinical features were also revealed, including correlations between lymphocyte and monocyte counts and between SARS-CoV-2 IgM or IgG and monocytes (reaching both *p* value and FDR significance) and other correlations reached nominal *p* value, but not FDR significance (**Figure 2B**), consistent with previous studies (*22*).

The relationship between CD63, CD32, C3aR and clinical disease indicated by ferritin levels in the blood suggested that the platelet activation potential of plasma from COVID-19 patients might provide additional insights into disease pathogenesis. Thromboembolic events are associated with poor outcomes and cardiopulmonary collapse in high-severity SARS-CoV-2 infected patients. Platelets not only contribute to normal thrombus formation but in case of hyperactivation can precipitate spontaneous clotting *in situ* (*29–32*). Thus, we examined the COVID-19 patients who experienced a clinically evident thrombosis during their hospital stay. We subdivided the samples to those coming from patients who did not yet have a clotting incident at the time of the blood draw (but went on to develop one later in their hospital stay) and the ones that already had a clotting event at or before the time of the draw. Plotting CD63 levels of healthy platelets activated by plasma from COVID-19 patients revealed a potential relationship between platelet activation capacity and future thrombotic event (**Figure 2D**). To further examine this potential relationship between platelet activation and thrombotic event, we compared CD63 induction by plasma from COVID-19 patients with future thrombotic events, patients with past thrombotic events, patients who never experience thrombotic complications and COVID-19 recovered patients (**Figure 2E**). Indeed, plasma obtained from patients prior to incident thrombotic events had the greatest capacity to induce CD63 on platelets from healthy donors compared to the other groups. Thus, platelet activation potential appears to be highest in COVID-19 patients prior to a clotting event.

### COVID-19 plasma-induced platelet activation is associated with markers of inflammation and cardiovascular disease

To begin to interrogate the soluble mediators that may underlie the platelet activation ability of COVID-19 patient plasma, we performed Proximity Extension Assays (PEA) using the O-link platform. This analysis interrogated 274 analytes in the blood with a focus on cardiovascular, inflammatory and organ damage related processes. We examined which of these circulating inflammatory mediators correlated with the ability of COVID-19 plasma to induce CD63 expression on platelets from healthy donors. Indeed, the concentration of numerous proteins in circulation correlated with the induction of CD63 on platelets (**Figure 3A**). The top proteins identified were mediators of inflammation, including IL-18, IL18BP, ADA, CCL15, NSF2, and proteins involved in vascular and heart pathology, including VEGFA, NPPC and PCSK9 (**Figure 3A**). There were also proteins suggesting neutrophil involvement such as NCF2, and potential indicators of nuclear content such as NBN, TOP2B, and EIF4EBP1 possibly consistent with release of neutrophil extracellular traps. It was notable that only positive correlations were revealed suggesting that, at least at this level of resolution, there were few if any counterregulatory pathways induced to maintain platelet quiescence. Individual positive correlations were examined more directly for CCL15, ADA, VEGFA and PCSK9 (**Figure 3B**). These findings further support the clinical metadata analysis described above (**Figure 2**) and establish the connection between increased platelet activation and circulating mediators of inflammation, neutrophil activity and/or indicators of cardiovascular disease and tissue damage in COVID-19.

**Figure 3.**
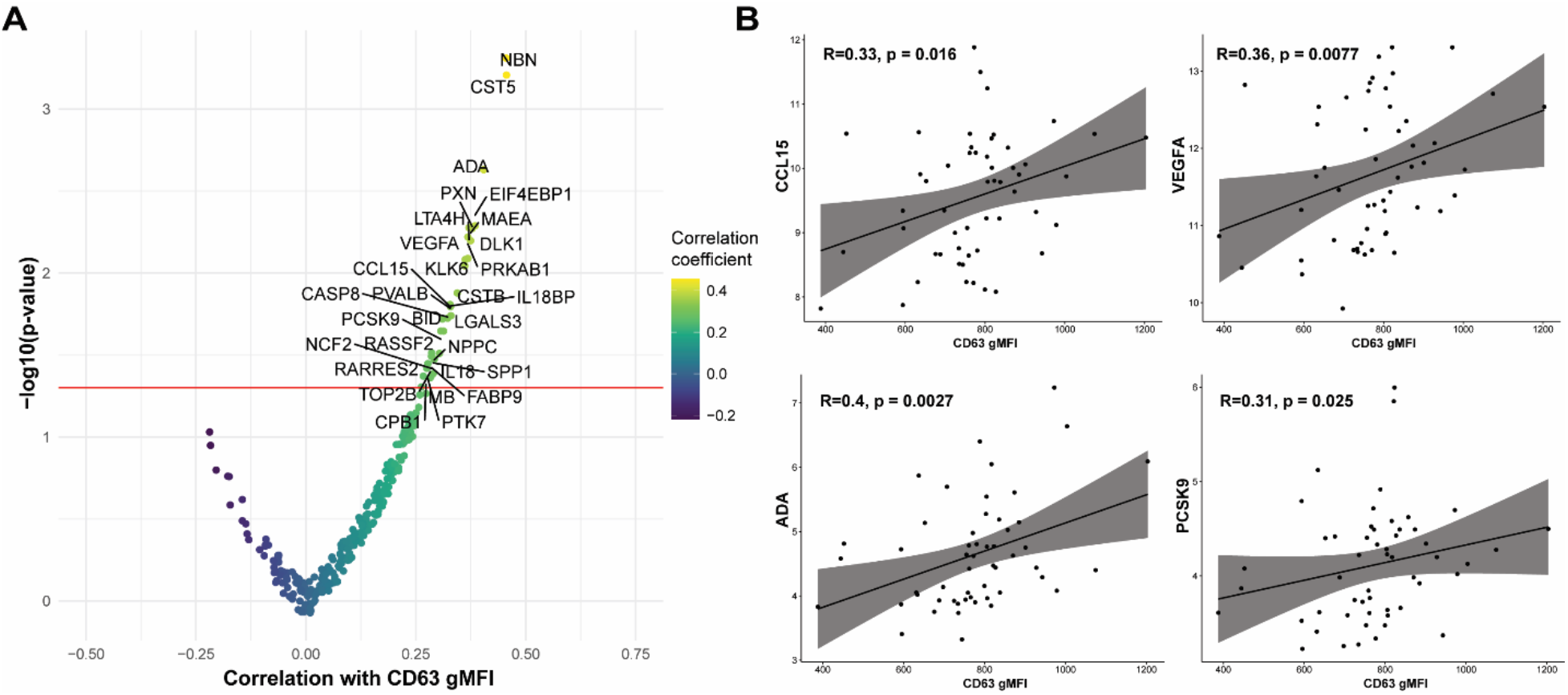
COVID-19 plasma induced platelet activation is associated with markers of inflammation and cardiovascular disease. **A)** Volcano plot with the x-axis representing the correlation coefficient of the different analytes with CD63 gMFI induction on platelets by COVID-19 plasma and the y-axis depicting the −log10 transformation of the corresponding *p*-value. **B)** Representative scatter plots for CD63 with CCL15, VEGFA, ADA and PCSK9.

### FcγRIIa activation and complement anaphylatoxins mediate platelet activation in COVID-19

The protein analysis described above suggest a connection between the platelet activation potential of COVID-19 plasma with tissue damage, cardiovascular pathways and COVID-19 associated inflammation. Infection with SARS-CoV-2 elicits a complex immune response, characterized by patterns of cellular and soluble inflammatory mediators that may differ considerably from patient to patient (*33–35*). How features of this inflammatory response relate to activation of platelets is unclear, though several possibilities exist. Specifically, immune complex formation and pro-inflammatory Fcγ structures have been implicated in the pathogenesis of severe COVID-19 (*36–38*) and platelets express FcγRIIa (*13, 16, 39*). In addition, platelets express complement receptors C3aR (*16*) and C5aR (*40*) and complement components, including anaphylatoxins, have role in the inflammatory cascade during SARS-CoV-2 infection (*19, 34, 35, 41*). Finally, IL-6 is often elevated and has been evaluated as a therapeutic target in COVID-19 (*42–44*) (*45, 46*). IL-6 may have a role in activating platelets in other settings, including through membrane-bound gp130 (*47*). As shown above, surface abundance of CD32 (FcγRIIa) and C3aR increase following platelet incubation with plasma derived from COVID-19 patients with a high inflammatory signature (**Figure 2B-C**). Thus, we hypothesized that Fc receptor signaling, IL-6 signaling and/or the signaling by the anaphylatoxins C3a and C5a, might be causally involved in platelet activation by COVID-19 plasma. To test this hypothesis, we examined the effect of blocking each of these pathways on the ability of plasma from COVID-19 patients to activate platelets (**Figure 4A** and **Supplementary Figure 3A**). Blocking each pathway individually inhibited platelet activation indicated by reduced induction of CD63, CD32, C3aR or CD62P. Blocking FcγRIIa had the strongest effect, followed by C5a neutralization whereas IL-6 and C3a blockade had the weakest effect and preferentially impacted CD62P and CD63, but not CD32 or C3aR (**Figure 4A**). However, blocking all 4 pathways simultaneously robustly decreased platelet activation indicated by all markers. This effect was more pronounced for samples from patients with higher ferritin (**Supplementary Figure 3B**). To evaluate how these distinct pathways might cooperate for platelet activation, we assessed different combinations of pathway blockade (**Figure 4B**). Although all pathways contributed to platelet activation indicated by CD63 induction, blockade of FcγRIIa appeared to have the most robust effect when combined with other blocking antibodies, especially in combination with anti-C5a and anti-C3a antibodies (**Figure 4B**). To further evaluate the role of FcγRIIa in platelet activation by COVID-19 plasma, we depleted IgG from the plasma samples prior to the platelet activation. IgG depletion reduced platelet activation by COVID-19 patient derived serum and this effect was accentuated further by neutralizing C5a (**Figure 4C**). Thus, these data indicate that the ability of plasma from COVID-19 patients to activate platelets occurs, at least partially, through IgG-mediated activation of FcγRIIa and this effect can be further augmented by signals from complement, including C5a.

**Figure 4.**
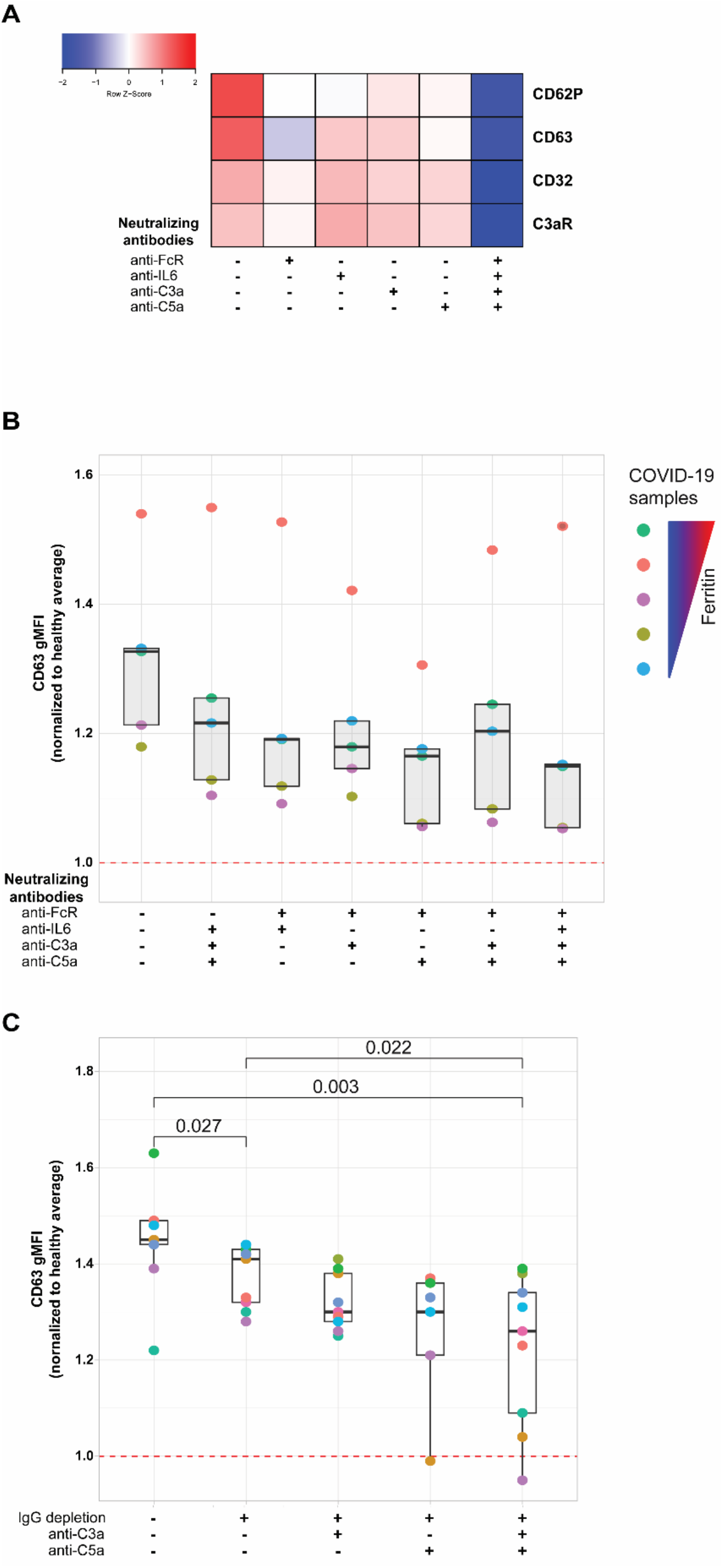
FcγRIIa activation and complement anaphylatoxins mediate platelet activation in COVID-19. **A)** Heatmap of gMFI expression of CD62P, CD63, CD32 and C3aR on the surface of control platelets incubated with COVID-19 plasma (n=10 patients) in the presence or absence of neutralizing antibodies to FcγRIIa, IL6, C3a and C5a, as indicated. **B)** Boxplots of gMFI expression of CD63 on the surface of control platelets incubated with COVID-19 plasma (n=5 patients, depicted with distinct colors) with different combinations of neutralizing antibodies to FcγRIIa, IL6, C3a and C5a, as indicated. **C)** Boxplots of gMFI expression of CD63 on the surface of control platelets incubated with COVID-19 plasma (n=9 patients, depicted with distinct colors) with different combinations of IgG depletion and neutralizing antibodies to C3a and C5a.

### Fostamatinib ameliorates the heightened activation of platelets induced by COVID-19 plasma

The FcγRIIa signals through recruitment and phosphorylation-mediated activation of Syk (*48, 49*). To confirm and extend the observations described above, we next investigated whether Syk inhibition impacted platelet activation by plasma from COVID-19 patients. Fostamatinib is a tyrosine kinase inhibitor that blocks the enzymatic activity of Syk and is clinically used for treatment of chronic immune thrombocytopenia (*50*). Syk phosphorylation was induced in as early as 1 minute following incubation of platelets from healthy donors with plasma from COVID-19 patients and increased further at 5 minutes compared to the effect of plasma from healthy control subjects (**Figure 5A**). We next assessed the impact of inhibiting Syk signaling on COVID-19 plasma mediated platelet activation. Addition of the fostamatinib active metabolite R406 to the platelet activation assay described above decreased induction of CD62P (**Figure 5B**) and CD63 (**Supplementary Figure 4**) on healthy platelets by plasma from COVID-19 patients. Addition of neutralizing antibodies to C3a and C5a did not further reduce platelet activation under these conditions suggesting a dominant role of Syk signaling in this setting. These data indicate a key role for antibody-mediated activation of platelets through FcγRIIa and possibly also complement activation. Whether this is the result of immune complexes, only IgG or also other isotypes that result in complement activation will require future studies. Nevertheless, these data suggest potential clinical utility of Syk inhibition or drugs that block complement signaling at least in a subset of patients.

**Figure 5.**
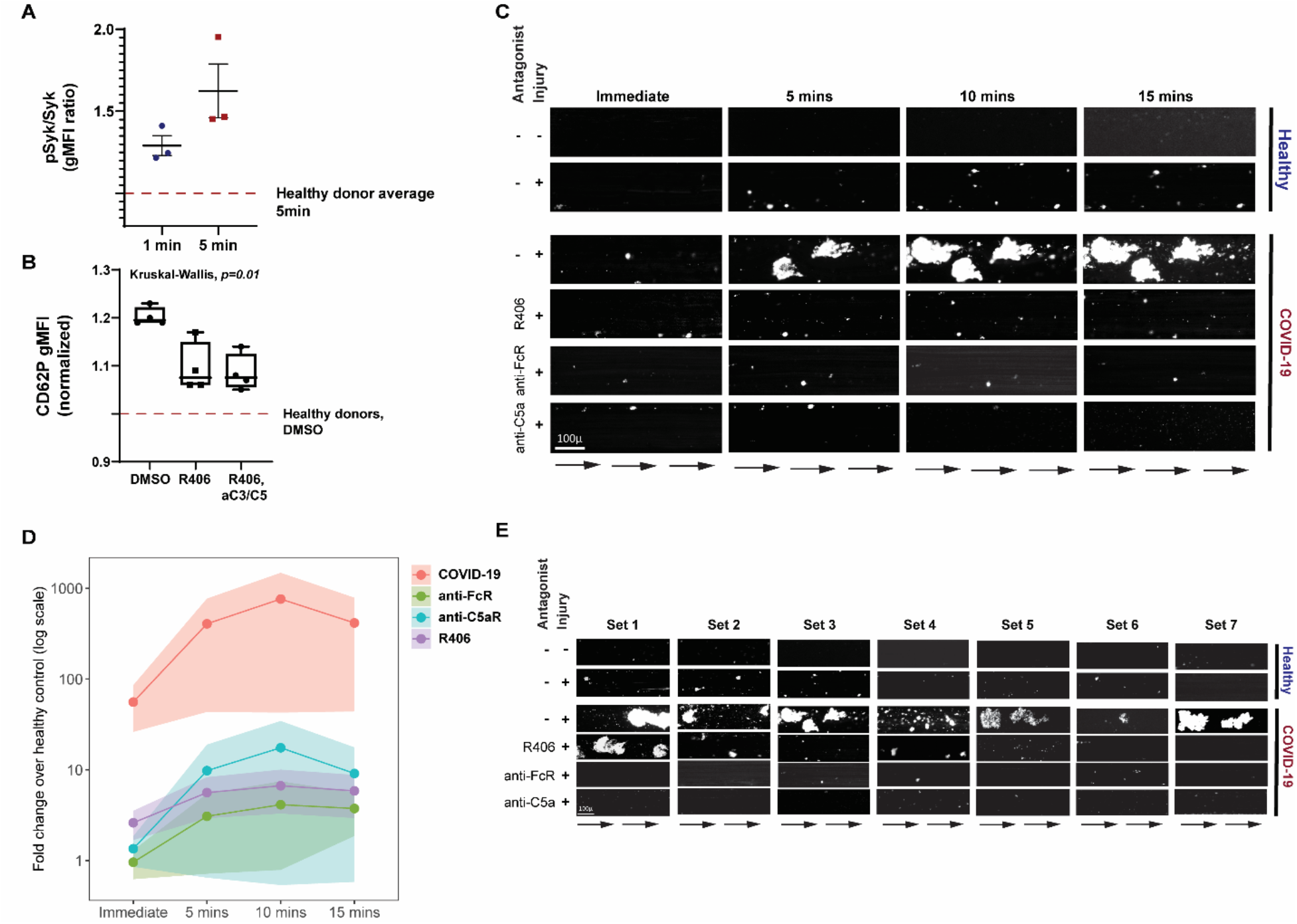
Fostamatinib ameliorates the heightened activation of platelets induced by COVID-19 plasma. **A)** Phospho-Syk and total Syk gMFI were measured with flow cytometry of control platelets incubated with COVID-19 plasma (n=3 patients) for 1 and 5 minutes. Incubation with healthy control plasma for 5 mins was used to normalize the data. **B)** Boxplots of gMFI expression of CD62P on the surface of control platelets incubated with COVID-19 plasma (n=4 patients) in the absence or presence of fostamatinib or fostamatinib and neutralizing antibodies to C3a and C5a. Kruskal-Wallis non-parametric testing was used to compare the groups and the *p*-value is depicted. **C)** Representative studies of a hematoporphyrin-induced photochemical injury model in an endothelial-lined microfluidic channel. Images show platelet adhesion (in white) immediately after infusion of isolated washed healthy donor platelets, and 5 min, 10 min and 15 min after infusion. Shown is a representative study where plasma from a severe COVID-19 patient was added to the platelets. In the samples indicated, the Syk inhibitor R406, an anti-FcγRIIa antibody, or an anti-C5a antibody were added prior to infusion. Direction of blood flow is indicated by arrows. Size bar indicates 100μ. **D)** Overall data analysis from studies with plasma derived from 7 patients with severe COVID-19. The Y-axis shows accumulated platelets in each microfluidic lane done immediately after the isolated platelet suspension was added and at immediate, 5, 10 and 15 minutes (log10 scale, fold change over healthy control plasma flown in uninjured channels). **E)** All 7 sets of experiments from D are shown at the 10-minute timepoint.

To further interrogate the prothrombotic potential of plasma from patients with COVID-19, we used a photochemical injury model in an endothelial-lined microfluidic channel. Induction of photochemical injury in this model results in activation of the endothelium with released von Willebrand factor (VWF) strands localized to the region of light exposure perfused with hematoporphyrin. We have previously described this approach in studies of platelet activation in the prothrombotic disorder of heparin-induced thrombocytopenia (*51*). Plasma from either normal control subjects or from patients with severe COVID-19 (1/10 final dilution) was added to calcein-labeled platelets isolated from healthy donors and perfused over injured endothelial cells. Although some platelet adhesion to the extruded VWF was observed over the subsequent 15 minutes for the channels where healthy plasma was added, this effect was markedly enhanced in those channels with added COVID-19 plasma (**Figures 5C-E**). Pre-incubation of platelets with either Syk inhibitor or antibodies to FcγRIIa, or C5a largely abrogated this enhanced platelet aggregation (**Figures 5C-E** and **Supplementary Table 2**). These data are consistent with a role for both FcγRIIa and complement activation of the platelets by the COVID-19 plasma and highlight the connection between antibody mediated platelet activation and the potential to initiate vascular thrombotic events. Moreover, the identification of these pathways dependent on Syk signaling suggests potential therapeutic opportunities.

## Discussion

Platelets act as a cellular connection between the immune system and hemostasis, integrating signals from different immune cell subsets, soluble inflammatory mediators and the complement cascade. As a result, platelets have the capacity to link signals from exuberant immune responses to thrombotic complications. Severe COVID-19 results in loss of immune homeostasis either due to a suboptimal, albeit persistent, immune response to SARS-CoV-2 or due to immune system hyperactivation that is disproportionate to what is necessary for efficient virus elimination (*52*). The data presented here indicate that plasma samples from COVID-19 patients with a high inflammatory index possess a robust ability to activate healthy platelets suggesting a link between infection-induced circulating mediators and potential thrombotic events. Indeed, induction of platelet activation was the highest in samples drawn prior to a clotting incident, an observation that could have future clinical utility. We were able to begin to identify pathways involved in this platelet activation revealing a key role for IgG mediated FcγRIIa signaling and complement, though other signals and pathways are also likely involved. Thus, our data define a key role for platelet activation potential of plasma from COVID-19 patients and link these observations to severity of the inflammatory state, specific immunological pathways, potential thrombotic risk, and a pathway with clinical targeting opportunities.

Cardiovascular disease risk factors, including hyperlipidemia, hypertension and diabetes, have been recognized since early in the pandemic as important determinants of COVID-19 outcomes (*23–25*). Our proteomics analysis revealed a correlation between mediators of cardiovascular health and platelet activation. Specifically, one of the most correlated proteins was VEGFA. Increased circulating VEGFA may point to an underlying strain to the cardiovascular system (*53*) in individuals with co-incident high levels of platelet activation. Correlation with other proteins of cardiovascular health, including NPPC (*54, 55*) and FABP9 (*56*), further support this hypothesis. Finally, among the highly correlated proteins was PCSK9 that has a central role not only in LDL metabolism (*57–59*) but has been shown to promote platelet activation (*60*) and can be found elevated in sepsis (*61*). As a result, our findings demonstrate a connection between platelet hyperactivation and elevated markers of cardiovascular dysregulation.

Complement activation and immune complexes that act through pro-inflammatory Fc structures have been linked to severe COVID-19 outcomes (*36, 38, 62, 63*). Inhibition of FcγRIIa signaling, either through blocking antibodies, IgG depletion or Syk inhibition, substantially reduced the platelet hyperactivation induced by COVID-19 plasma. Fostamatinib has an FDA-approved indication as first-line treatment for chronic ITP (*50*). This drug has also been trialed in rheumatoid arthritis, albeit with modest effects (*64*). In this report, Syk inhibition using fostamatinib robustly blunted *in vitro* platelet activation induced by COVID-19 plasma. Not only do these data identify a potential mechanism of platelet activation in COVID-19 via antibody and/or immune complex mediated platelet Fc receptor signaling, but they also point to a therapeutic intervention opportunity in COVID-19 patients, especially those with a high risk for thrombotic complications. Indeed, fostamatinib is currently being tested in a clinical trial in hospitalized COVID-19 patients (ClinicalTrials.gov Identifier: NCT04579393). A preliminary report posted by the manufacturer of fostamatinib suggested that Syk inhibition might protect against COVID-19 induced hypoxia. Although speculative at this stage, this information might indicate that Syk is required for the formation of pulmonary microthrombi reported in patients with COVID-19 induced hypoxia (*31*). In addition to Syk inhibition, inhibition of the Fc-mediated and/or complement-mediated platelet activation could be achieved through other therapeutic modalities. For example, Fc-mediated platelet stimulation could be tempered by plasmapheresis of immune complexes from COVID-19 patients (ClinicalTrials.gov Identifier: NCT04374539). In addition, it may also be relevant that of the two complement anaphylatoxin receptors tested, C5a neutralization had the largest additive effect to Fc blockade reducing platelet hyperactivation. Blocking the C5a-C5aR axis with monoclonal antibodies against C5aR reduced the activation of human myeloid cells and lung injury in a human C5aR knock-in mouse model (*35*). Eculizumab, a monoclonal antibody that inhibits the cleavage of C5 to C5a and C5b and previously approved for treatment of paroxysmal nocturnal hematuria (*65*) and atypical hemolytic uremic syndrome (*66*), is also currently in a clinical trial to investigate its efficacy in severe COVID-19 (ClinicalTrials.gov Identifier: NCT04355494). Other potential complement pathway inhibitors also exist and are under investigation in COVID-19 patients. The current work may provide a deeper understanding of how these interventions function to prevent thromboembolic events, suggest biomarkers of drug efficacy and potential drug combinations that may be of benefit in some patients. Thus, our data identify the FcγRIIa-Syk and the C5a-C5aR axes as key mediators of platelet hyperactivity in COVID-19 and highlight the therapeutic potential of targeting these mechanisms in COVID-19 patients, especially those with high-thrombotic risk.

Finally, our findings provide a framework for additional platelet-focused studies. They establish the role of thromboinflammation in COVID-19, support the role of cardiovascular disequilibrium in platelet dysfunction and indicate that there are plasma soluble factors that drive platelet hyperactivation. Moreover, given the recent concerns about very rare venous thrombosis and thrombocytopenia following adenovirus-based SARS-CoV-2 vaccination (*67–70*), the studies described here might suggest approaches to evaluate underlying mechanisms for these events. Indeed, although studies are necessary to examine the beneficial potential of restoring platelet function in COVID-19 patient outcomes, our studies highlight putative therapeutic candidates to address platelet-driven clotting complications of COVID-19.

## Materials and Methods

### Patients, subjects, and clinical data collection

Patients admitted to the Hospital of the University of Pennsylvania with a SARS-CoV-2 positive result were screened and approached for informed consent within 3 days of hospitalization (COVID-19 inpatient group). Health care workers were recruited at the Hospital of the University of Pennsylvania and received both a PCR test to assess for active infection and serologic testing for antibodies against SARS-CoV-2; all individuals included in this study were serologically convalescent and PCR negative (COVID-19 convalescent group). Healthy donors were recruited through word of mouth. Peripheral blood was collected from all subjects. For inpatient cases, clinical data were abstracted from the electronic medical record into standardized case report forms. APACHE III scoring was based on data collected in the first 24 hours of ICU admission or the first 24 hours of hospital admission for subjects who remained in an inpatient unit. Comorbidities including prior history of clotting (deep vein thrombosis (DVT), pulmonary embolism (PE), cerebrovascular accident (CVA), myocardial infarction (MI), or other thrombus) were collected prospectively based on the EMR. Cardiovascular disease (CVD) risk factors were considered present if patients had any of the following: diabetes, hypertension, hyperlipidemia, peripheral arterial disease, cerebrovascular disease, or known coronary artery disease. In-hospital clotting events were determined by EMR chart review and requiring a documented and date-stamped DVT, PE, CVA, MI, or other thrombus by duplex, echocardiogram, or contrast-enhanced imaging. Anticoagulation (AC) treatment at the time of research blood collection was recorded; “intermediate” dose AC was equivalent to enoxaparin 0.5 mg/kg subcutaneously twice daily in contrast to “prophylactic” dose enoxaparin of 0.5 mg/kg once daily. Clinical laboratory data were collected from the date closest to the date of research blood collection. CVD risk was defined at the presence of pre-existing hypertension, hyperlipidemia and/or diabetes mellitus. All participants or their surrogates provided informed consent in accordance with protocols approved by the regional ethical research boards and the Declaration of Helsinki.

### Sample processing, platelet isolation and activation

Peripheral blood was collected into sodium heparin tubes (BD, Cat#367874). For directly *ex vivo* assays, whole blood was diluted 10-fold with Tyrode’s buffer and then used for stimulation assays with TRAP or vehicle in the presence of 50mM CaCl2. TRAP was added at a final concentration of 35uM for 15mins at 37°C. The activation was terminated by adding PBS/4% PFA for 20 mins. For *in vitro* activation assays of control platelets with COVID-19 plasma, healthy platelets were first isolated. Whole blood collected in citrate tubes was spun at 125g for 15 minutes and the supernatant was collected to isolate PRP (platelet rich plasma). PRP was spun at 330g for 10 minutes to collect pelleted platelets that were resuspended in 1x Tyrode’s Buffer. Platelets were subsequently activated on a 96-well plate with 50uL of sodium heparin isolated plasma derived from COVID-19 patients or healthy volunteers.

### Neutralization and inhibition assays

Neutralizing monoclonal antibodies against C3a (Biolegend, cat # 518105), C5a (R&D, cat # MAB2037-SP), IL-6 (Biolegend, cat # 501101) and CD16/32 (Biolegend, cat # 101302) were used for the neutralization assays. IgG depletion from plasma was performed using the Albumin IgG Depletion Spintrap from Millipore-Sigma (cat # GE28-9480-20) based on the manufacturer’s instructions. The active metabolite of fostamatinib R406 was purchased from Selleckchem (cat # S1533), dissolved in DMSO and used at a final concentration of 5μM.

### Flow Cytometry and antibody clones

Antibodies used for staining of whole blood and platelets:

|  |  |  |
| --- | --- | --- |
| BB700 Mouse Anti-Human CD32 | BD | Cat # 745929 |
| APC Mouse Anti-Human CD42b | BD | Cat # 555524 |
| PE Mouse Anti-Human CD62P | BD | Cat # 555524 |
| PE/Cyanine7 anti-human C3AR Antibody | Biolegend | Cat # 345808 |
| BV421 anti-human CD63 Antibody | Biolegend | Cat # 353030 |

Samples were acquired on a 4 laser BD FACS LSR. Standardized SPHERO rainbow beads (Spherotech, Cat#RFP-30-5A) were used to track and adjust PMTs over time. UltraComp eBeads (ThermoFisher, Cat#01-2222-42) were used for compensation. Up to 1×10^5^ platelets were acquired per each sample.

### Proximity Extension Assays and SARS-CoV-2 Serologic Testing

Proximity Extension Assays using plasma derived from COVID-19 patients were performed using the commercially available Olink protein biomarker platform. Specifically, three Olink Target 96 panels were used: Olink Target 96 Cardiovascular III, Olink Target 96 Inflammation and Olink Target 96 Organ Damage panel. SARS-CoV2-RBG IgG and IgM measurements were performed by enzyme-linked immunosorbent assays (ELISA) as previously described (*71*).

### Platelet aggregation on photochemically-injured endothelium in a microfluidic system

6 × 10^6^ cells/channel of human umbilical vein endothelial cells (HUVECs, ATCC-PCS-100-013) were seeded into the fibronectin (50 μg/mL, Sigma-Aldrich cat. # F0895) coated channels of a 48-well microfluidic plate (Bioflux, Fluxion Biosciences) and then injured by flowing a 50ug/mL solution of hematoporphyrin (Sigma-Aldrich) with exposure to blue light using the HXP-120 C light source with 475-nm excitation and 530-nm emission filters as previously described (*51*). 200μL of platelet suspension (2×10^6^ platelets in HBSS^Ca-Mg-^) from healthy donors was labelled with calcein-AM (2 μg/mL final concentration, ThermoFisher Scientific Cat # C3100MP) for 15 mins and then R406, anti-FcγRIIa or anti-C5a were added into the respective tube for a 15-minute incubation. 20μl of either healthy control or COVID-19 patient plasma (1:10 final dilution) was added just prior to being flowed through the channel at 10 dynes/cm^2^. Platelet accumulation in the injured endothelium field was captured by Zeiss Axio Observer Z1 inverted microscope using Montage Fluxion software and analyzed using ImageJ as described (*51*).

### Correlation plot and statistics

Pairwise correlations between variables were calculated and visualized as a correlogram using the R function *corrplot* (Figure 2B) displaying the positive correlations in red and negative correlations in blue. Spearman p-value significance levels were shown.

Due to the heterogeneity of clinical and flow cytometric data, non-parametric tests of association were preferentially used throughout this study unless otherwise specified. Correlation coefficients between ordered features (including discrete ordinal, continuous scale, or a mixture of the two) were quantified by the Spearman rank correlation coefficient and significance was assessed by the corresponding non-parametric methods (null hypothesis: ρ = 0). Tests of association between mixed continuous versus non-ordered categorical variables were performed by Mann-Whitney U test (for n = 2 categories) or by Kruskal-Wallis test (for n > 2 categories). All tests were performed two-sided, using a nominal significance threshold of P < 0.05 unless otherwise specified. When appropriate to adjust for multiple hypothesis testing, false discovery rate (FDR) correction was performed by the Benjamini-Hochberg procedure at the FDR < 0.05 significance threshold unless otherwise specified.

Other statistical analysis was performed using Prism software (GraphPad). Other details, if any, for each experiment are provided within the relevant figure legends.

## List of supplementary materials

**Supplementary Figure 1.** Cumulative data for CD62P surface expression of *ex vivo* isolated platelets at baseline (basal), after TRAP activation (TRAP) and their ratio for hospitalized COVID-19 patients that experienced a clinical thrombosis or not. Mann-Whitney non-parametric testing was used and the *p*-values are depicted.

**Supplementary Figure 2.** Gating strategy for isolated control platelets incubated with COVID-19 plasma.

**Supplementary Figure 3. A)** Violin plots of gMFI expression of CD62P, CD63, CD32 and C3aR on the surface of control platelets incubated with COVID-19 plasma (n=10 patients) in the presence or absence of neutralizing antibodies to FcγRIIa, IL6, C3a and C5a, as indicated. Kruskal-Wallis non-parametric testing was used to compare the groups and the *p*-values are depicted. **B)** Same samples and conditions depicted in (A) but categorized based on the corresponding patient’s ferritin levels. Ferritin – low: <1000ng/mL; Ferritin – medium: 1000-2000ng/mL; Ferritin – high >2000ng/mL

**Supplementary Figure 4.** Boxplots of gMFI expression of CD63 on the surface of control platelets incubated with COVID-19 plasma (n=4 patients) in the absence or presence of fostamatinib or fostamatinib and neutralizing antibodies to C3a and C5a.

**Supplementary Table 1.** Clinical information of the COVID-19 patients evaluated.

**Supplementary Table 2**. Inhibition of platelet aggregation in hematoporphyrin-induced photochemical injury model in an endothelial-lined microfluidic channel. Analysis of relative fluorescence intensity of platelet aggregation in the hematoporphyrin-induced photochemical injured endothelial-lined microfluidic channel at different times after infusion of platelets in plasma (n=7) from a severe COVID-19 patient: immediate, 5 min, 10 min and 15 min. Data is expressed as fold increase of platelet accumulation with patient plasma over healthy donor plasma. P values were calculated using Dunnett’s multiple comparisons test, n=7.

## Acknowledgements

We would like to sincerely thank patients and blood donors, their families and surrogates, and medical personnel. The UPenn COVID Processing Unit comprises individuals from diverse laboratories at the University of Pennsylvania who volunteered time and effort to enable study of COVID-19 patients during the pandemic: Sharon Adamski, Zahidul Alam, Mary M. Addison, Katelyn T. Byrne, Aditi Chandra, Hélène C. Descamps, Nicholas Han, Yaroslav Kaminskiy, Shane C. Kammerman, Justin Kim, Jacob T. Hamilton, Nune Markosyan, Julia Han Noll, Dalia K. Omran, Eric Perkey, Elizabeth M. Prager, Dana Pueschl, Austin Rennels, Jennifer B. Shah, Jake S. Shilan, Nils Wilhausen, Ashley N. Vanderbeck. All are affiliated with the University of Pennsylvania Perelman School of Medicine.

## Author contributions

SAA, CSA and EJW conceived the study. SAA, AmS, HMG, RRG, DM, MAH and SG carried out experiments. SAA, HG, AEB, ARG, CA, JEW, ZC, YJH, AP, OK, JD, ARW, CAGI, DD, IF, ACH, LAV, JPR and NM were involved in clinical recruitment, sample allocation, processing and acquisition. All authors participated in data analysis and interpretation. SAA, AmS, AaS, LR, LZ, MP, CSA and EJW wrote the manuscript. All authors reviewed the manuscript.

## Data and materials availability

All data associated with this study are available in the main text or the supplementary materials.

