## Supplementary_materials for "Signaling through FcγRIIA and the C5a-C5aR pathway mediates platelet hyperactivation in COVID-19"

**Supplementary Figure 1.**

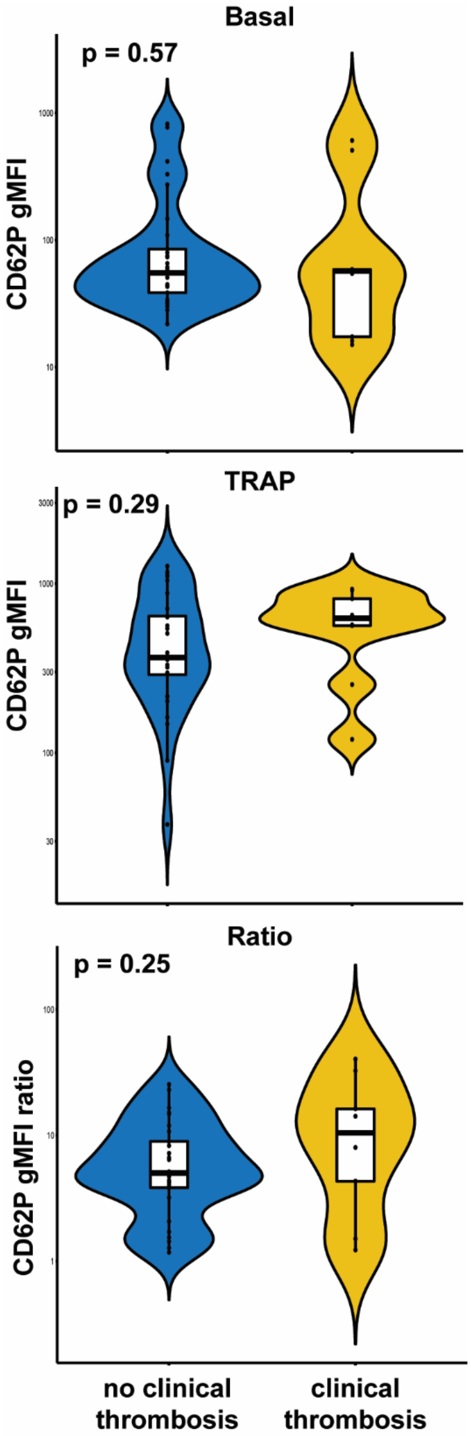


**Supplementary Figure 1.** Cumulative data for CD62P surface expression of *ex vivo* isolated platelets at baseline (basal), after TRAP activation (TRAP) and their ratio for hospitalized COVID-19 patients that experienced a clinical thrombosis or not. Mann-Whitney non-parametric testing was used and the *p*-values are depicted.

**Supplementary Figure 2.**

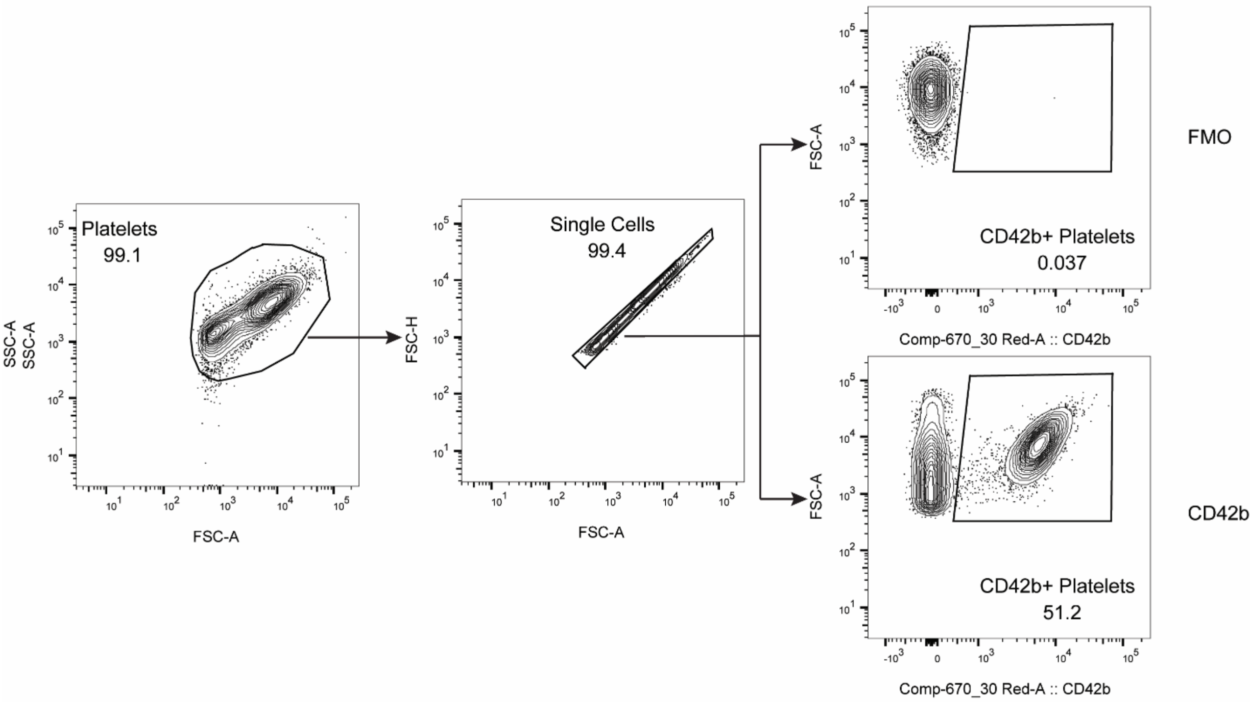


**Supplementary Figure 2.** Gating strategy for isolated control platelets incubated with COVID-19 plasma.

**Supplementary Figure 3.**

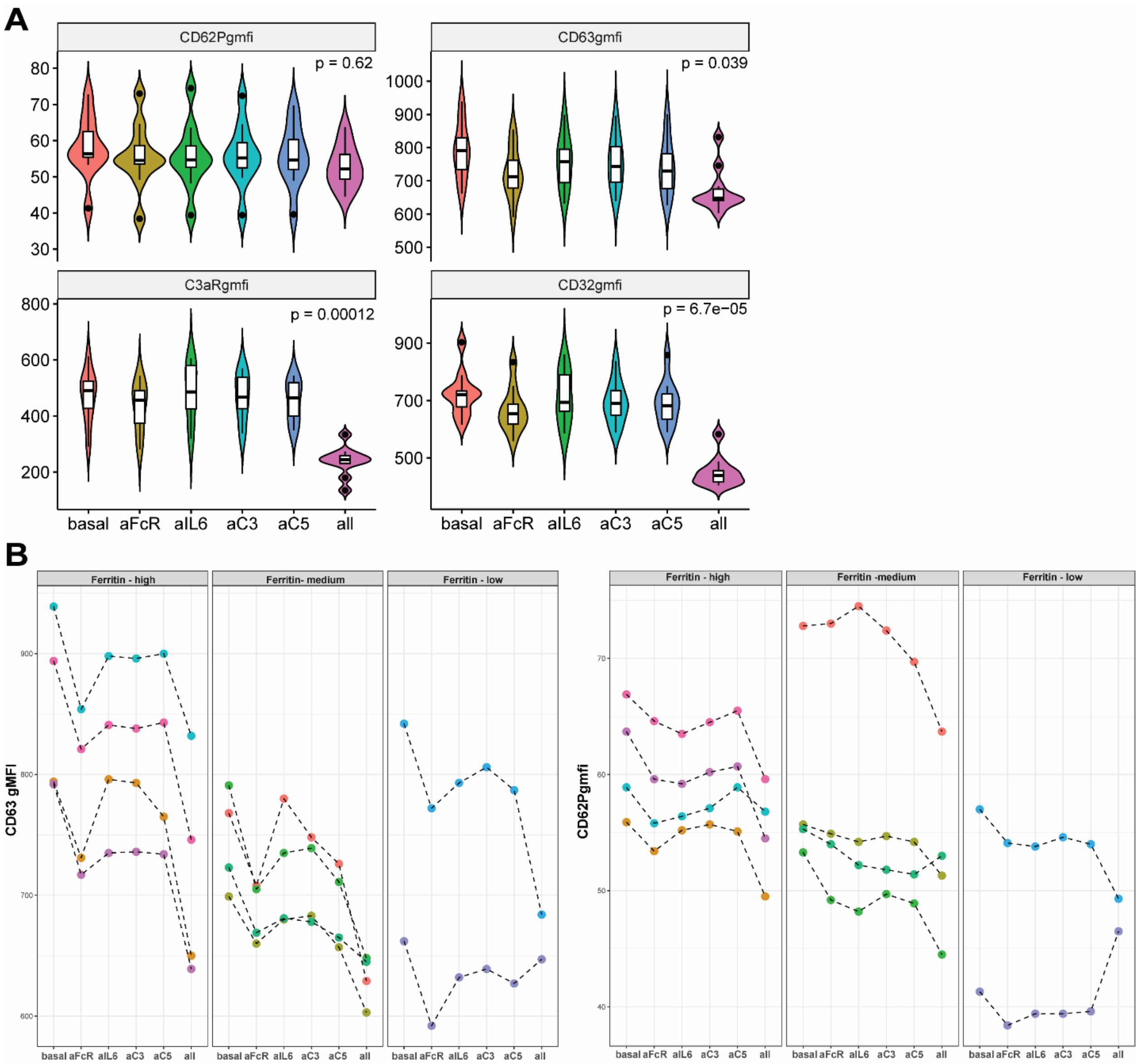


**Supplementary Figure 3. A)** Violin plots of gMFI expression of CD62P, CD63, CD32 and C3aR on the surface of control platelets incubated with COVID-19 plasma (n=10 patients) in the presence or absence of neutralizing antibodies to FcγRIIa, IL6, C3a and C5a, as indicated. Kruskal-Wallis non-parametric testing was used to compare the groups and the *p*-values are depicted. **B)** Same samples and conditions depicted in (A) but categorized based on the corresponding patient's ferritin levels. Ferritin – low: <1000ng/mL; Ferritin – medium: 1000-2000ng/mL; Ferritin – high >2000ng/mL

**Supplementary Figure 4.**

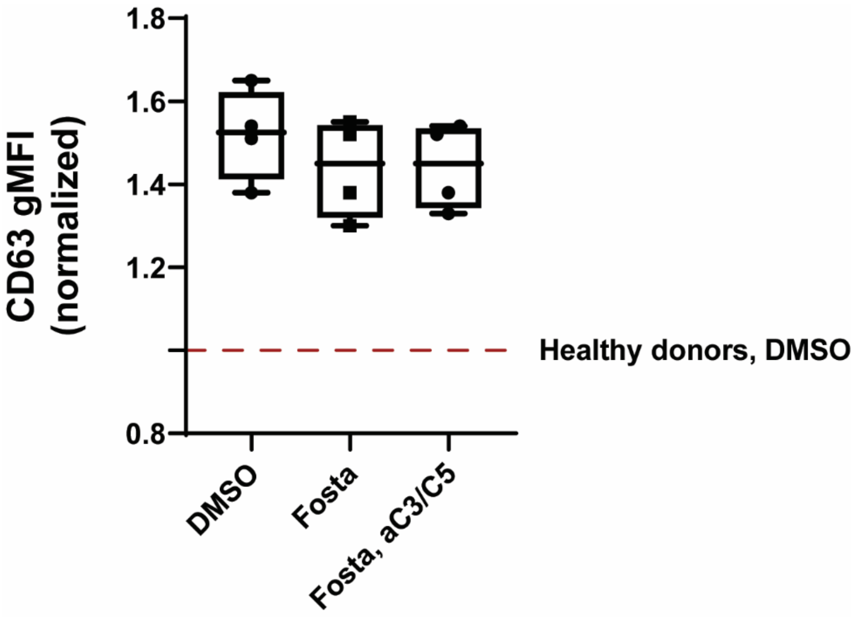


**Supplementary Figure 4.** Boxplots of gMFI expression of CD63 on the surface of control platelets incubated with COVID-19 plasma (n=4 patients) in the absence or presence of fostamatinib or fostamatinib and neutralizing antibodies to C3a and C5a.

**Supplementary Table 1.**

| | COVID-19 patients with platelets assayed *ex vivo* (33 unique patients, 38 samples) | COVID-19 patients with plasma assayed *in vitro* (47 unique patients, 64 samples) |
|---|---|---|
| Age (mean, SEM) | 60.2 (2.94) | 58.7 (2.14) |
| Gender (male, %) | 48% | 62% |
| BMI (mean, SEM) | 34 (2.2) | 32.7 (1.67) |
| Platelet count (mean, SEM) | 261.9 (17.8) | 246 (12.6) |
| WBC count (mean, SEM) | 7.9 (0.6) | 9.3 (1.0) |
| D-dimer (mean, SEM) | 4.37 (1.64) | 7.04 (2.8) |
| Ferritin (mean, SEM) | 1350.3 (363) | 1650.1 (313.8) |
| LDH (mean, SEM) | 347.2 (22.9) | 404.5 (33.25) |
| Hs-CRP (mean, SEM) | 91.8 (10.9) | 81.53 (12.57) |
| CVD risk (%) | 82% | 87% |
| APACHEIII (mean, SEM) | 61.6 (3.9) | 63.58 (4.27) |
| Enrollment NIH Disease Severity Score (mean, SEM) | 3.4 (0.18) | 3.28 (0.15) |
| Incident thrombosis (%) | 9% | 34% |
| SARS-CoV-2 IgM (ug/mL, mean, SEM) | 10.47 (3.73) | 10.53 (4.48) |
| SARS-CoV-2 IgG (ug/mL, mean, SEM) | 41.1 (8.76) | 47.9 (10.65) |

**Supplementary Table 1.** Clinical information of the COVID-19 patients evaluated.

**Supplementary Table 2**.

| | Condition | Fold-change (mean ± SEM) | P value vs. No drug |
|---|---|---|---|
| **Immediate** | --- | 55.85 ± 29.63 | |
| | R406 | 2.61 ± 0.92 | 0.0255 |
| | Anti-FcR | 0.96 ± 0.34 | 0.0245 |
| | Anti-C5a | 1.35 ± 0.50 | 0.0275 |
| **5 mins** | --- | 406.65 ± 363.08 | |
| | R406 | 5.613 ± 2.68 | 0.0255 |
| | Anti-FcR | 3.08 ± 2.36 | 0.0245 |
| | Anti-C5a | 9.79 ± 9.14 | 0.0275 |
| **10 mins** | --- | 762.60 ± 719.81 | |
| | R406 | 6.68± 3.373 | 0.0255 |
| | Anti-FcR | 4.13 ± 3.34 | 0.0245 |
| | Anti-C5a | 17.52 ± 16.98 | 0.0275 |
| **15 mins** | --- | 415.88 ± 371.80 | |
| | R406 | 5.86 ± 2.92 | 0.0255 |
| | Anti-FcR | 3.75 ± 1.88 | 0.0245 |
| | Anti-C5a | 9.13 ± 8.54 | 0.0275 |

**Supplementary Table 2**. Inhibition of platelet aggregation in hematoporphyrin-induced photochemical injury model in an endothelial-lined microfluidic channel. Analysis of relative fluorescence intensity of platelet aggregation in the hematoporphyrin-induced photochemical injured endothelial-lined microfluidic channel at different times after infusion of platelets in plasma (n=7) from a severe COVID-19 patient: immediate, 5 min, 10 min and 15 min. Data is expressed as fold increase of platelet accumulation with patient plasma over healthy donor plasma. P values were calculated using Dunnett’s multiple comparisons test, n=7.
